# Essential role of an acidic endosomal environment in JAK3-mediated signal transduction

**DOI:** 10.1101/2023.09.07.556780

**Authors:** Yuji Amano, Nobuyuki Tanaka, Toshikazu Takeshita, Katsuhiko Kojima

**Affiliations:** Department of Microbiology and Immunology, Shinshu University School of Medicine, Matsumoto, Japan; Division of Tumor Immunobiology, Miyagi Cancer Center Research Institute, Natori, Sendai, Japan; Division of Tumor Immunobiology, Tohoku University Graduate School of Medicine, Sendai, Japan

**Keywords:** Cytokine, signal transduction, Janus kinase, endosome, spatial regulation, chloroquine, anti-inflammatory drug

## Abstract

The endosome, an acidic organelle, is a platform for endocytic trafficking and signal transduction. Chloroquine, an intercellular acidic vesicle neutralizer (IAVN), has been used to treat autoimmune diseases. Its immunosuppressive mechanism remains unclear, although several modes of action attributable to lysosomal dysfunction have been postulated. This study found that IAVNs, including chloroquine, can selectively interfere with common cytokine receptor γ chain (γc) cytokine-induced signal transduction. Our data show that JAK3 targets endosomal membranes in an acidic pH-dependent manner, which is essential for γc-cytokine-induced functional signal complex formation and signal activation. Among IAVNs, Monensin and similar ionophores were particularly effective in suppressing γc-cytokine-induced cell proliferation in vitro. Monensin exerted therapeutic effects in a collagen-induced arthritis mouse model by interfering with T helper 17 cells. Our findings help further understand the immunosuppressive mechanism of IAVNs and unveil the spatial regulation of JAK3-mediated signal transduction via endosomes.

## Introduction

Following ligand binding at the plasma membrane, cytokine and growth factor receptors are endocytosed and transported to lysosomes for degradation via endosomal trafficking. This process plays a role in receptor downregulation and attenuation of receptor-mediated signal transduction^1, 2^. Furthermore, in some cases, endocytic trafficking of signaling receptors can generate specific signals that depend on the endosomal environment ^3–6^. Endosomes have unique membrane surface and lumen properties that permit the recruitment of endosomal scaffold and adaptor proteins. The enrichment of phosphatidylinositol 3-phosphate (PtdIns3P) is a characteristic feature of the endosomal membrane, which allows the assembly of complexes involving molecules with FYVE and PX domains ^7–10^. An acidic environment in the endosomal lumen is also a critical aspect that regulates pH-dependent binding between the FYVE domain and PtdIns3P ^11, 12^.

Chloroquine (CQ) is a diprotic base that can accumulate in acidic organelles such as endosomes and lysosomes. Moreover, it causes an increased pH in acidic vesicle lumens by trapping H^+^ ions ^13^. CQ was originally identified as an anti-malarial drug and later discovered to have a therapeutic effect on rheumatoid arthritis (RA). Thereafter, CQ and its derivative hydroxychloroquine (HCQ) have been used as immunosuppressants in patients with RA, systemic lupus erythematosus, and other autoimmune diseases ^14, 15^. It is well known that endosomal acidification is essential for all living cells to maintain cellular homeostasis, including lysosomal proteolysis, autophagy, and vesicular trafficking ^16–18^. CQ has a well-established and excellent safety profile when used at the recommended doses, although CQ is extensively distributed throughout the entire body after administration ^14, 15^. This evidence implies that there is a mechanism influencing the immune system involving neutralization of acidic organelles more sensitive than other physiological functions. The mechanism controlling the immunomodulatory effect of CQ remains uncertain, though various modes of action have been postulated, including inhibition of major histocompatibility complex (MHC) class II-mediated autoantigen presentation and interfering with Toll-like receptor (TLR) 7 and TLR9-mediated signal transduction ^14, 19–22^.

The Janus kinases (JAKs), JAK1, JAK2, JAK3, and TYK2, form a family of non-receptor tyrosine kinases that principally mediate cytokine-induced signal transduction ^23^. JAKs are involved in the growth, survival, development, and differentiation of various cell types, but also play particularly important roles in immune cells, which make them an attractive molecular target for the treatment of allergic and autoimmune diseases ^24, 25^. Among JAKs, JAK3 exclusively functions in immune cells. Like the common cytokine receptor γ chain (γc), JAK3 is predominantly expressed in hematopoietic cells and plays an essential role in signal activation of the γc-cytokines, interleukin (IL)-2, IL-4, IL-7, IL-9, IL-15, and IL-21, which are indispensable for lymphocyte survival, proliferation, and differentiation ^26, 27^.

Cytokine binding to the extracellular domain of their respective receptors leads to activation of the JAK/STAT pathway, essential for maintaining immune homeostasis and inflammatory responses ^23^. This step induces receptor dimerization and a structural change in the cytoplasmic domain of the receptor subunits, resulting in the activation of JAKs ^28–30^. This is followed by the phosphorylation of downstream signaling molecules, such as the Signal transducer and activator of the transcription (STAT) family of transcription factors ^31, 32^. In some cytokine receptor subunits, a cytokine-induced structural change is also necessary for interaction with JAKs ^33, 34^. Interactions between JAKs and receptor subunits are critical for cytokine-induced signal transduction, but the molecular details of these interactions are unclear.

This report demonstrates that intercellular acidic vesicle neutralizers (IAVNs), including CQ, can suppress γc-cytokine-induced phosphorylation of STATs. We further show that JAK3 targets the endosomal membrane in an endosomal acidic pH-dependent manner, and this is necessary for the assembly of functional signaling complexes and JAK3-mediated signal transduction.

## Results

### Neutralization of intercellular vesicles causes inhibition of γc-cytokine-induced signal transduction

To examine whether CQ treatment affects the cytokine-mediated activation of STATs, we stimulated human peripheral blood mononuclear cells (PBMCs) with various cytokines in the presence or absence of CQ. The results showed that IL-2, IL-7, IL-15, and granulocyte macrophage colony stimulating factor (GM-CSF)-induced STAT phosphorylation were inhibited by CQ treatment (Fig. 1A–D). In contrast, CQ did not affect IL-6, IL-10, interferon (IFN)α or IFNγ-induced STAT phosphorylation (Fig. 1E–H). Similar results were observed with Monensin and ammonium chloride, which can neutralize intercellular acidic vesicles like CQ, suggesting that certain cytokine-induced signal transduction requires intercellular vesicles with an acidic environment (Fig. 1A–H). Notably, γc-cytokine-induced STAT phosphorylation was consistently inhibited by IAVNs, while other cytokine signals, except GM-CSF, were not affected. To further verify the effect of IAVNs on γc-cytokine-induced signal activation, we established Ba/F cells that can activate and proliferate by both IL-3 and γc-cytokines: IL-2, IL-4, or IL-7 (Supplementary Fig. 1A–C) ^35–37^. Cytokine-induced cell proliferation was compared between IL-3 and γc-cytokines in the presence of IAVNs. Consistent with the effect on STAT phosphorylation, IAVNs suppressed all three γc-cytokine-dependent cell proliferation (Fig. 1I–N). Although the IAVNs also inhibited the IL-3-dependent cell proliferation, EC50s of CQ and ammonium chloride were approximately one-fifth to half the concentrations compared with IL-3-induced cell proliferation (Fig. L–N). Remarkably, Monensin most potently inhibited γc-cytokine-induced cell proliferation compared to the relatively modest effects of CQ and ammonium chloride; EC50 79.3 nM (IL-2), 4.4 nM (IL-4), 2.6 nM (IL-7). Because Monensin is a carrier-type ionophore that acts as a proton/cation antiporter ^38^, we also tested other proton/cation antiporters, Nigericin and Salinomycin^13, 39^. These two agents also inhibited IL-2, IL-4, and IL-7 -induced cell proliferation while modest inhibition of IL-3-dependent growth (Supplementary Fig. 1D–I). These results suggest that IAVNs potentially inhibit the γc-cytokine-dependent cell proliferation in a relatively selective manner.

**Figure 1.**
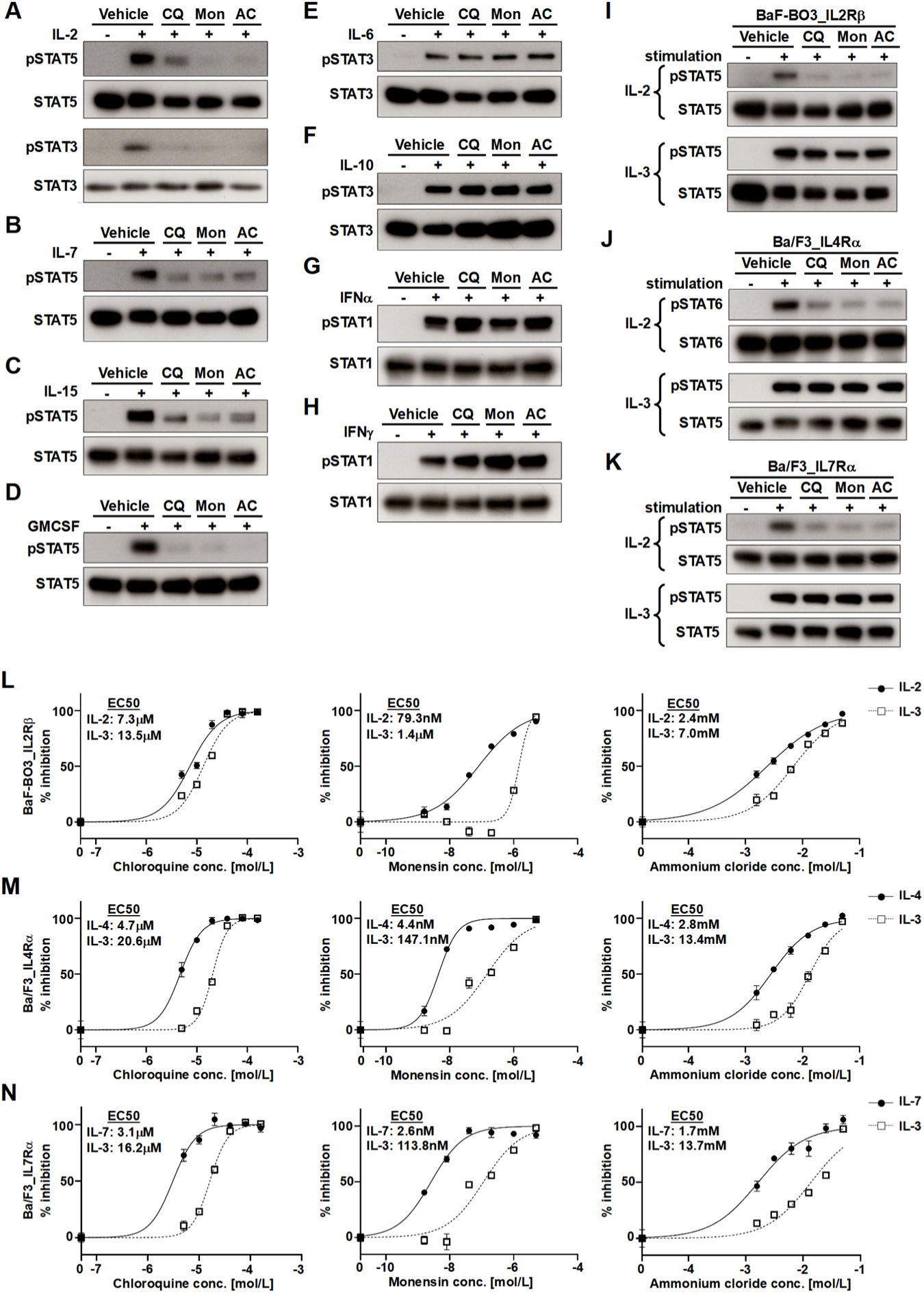
Intercellular acidic vesicle neutralizers (IAVNs) interfere with γc-cytokine-induced signal transduction. (A–H) The effects of IAVNs on various cytokine-induced STAT phosphorylation in peripheral blood mononuclear cells (PBMCs). Freshly isolated PBMCs were pre-incubated with 80 μM chloroquine (CQ), 1 μM Monensin (Mon), or 50 mM ammonium chloride (AC) for 1 hour, then stimulated with (A) interleukin (IL)-2, (B) IL-7, (C) IL-15, (D) granulocyte-macrophage colony-stimulating factor (GM-CSF), (E) IL-6, (F) IL-10, (G) interferon (IFN) α, or (H) IFNγ for 30 minutes. (I–K) Selective suppressive effects of IAVNs on γc-cytokine-induced STATs phosphorylation in transfected Ba/F cells. (I) BaF-B03_IL2Rβ, (J) Ba/F3_IL4Rα, and (K) Ba/F3_IL7Rα cells were pre-incubated with 80 μM CQ, 1 μM Mon, or 50 mM AC for one hour, then stimulated with IL-3 and IL-2, IL-4, or IL-7, respectively, for 10 minutes. (L–N) Selective inhibitory effects of IAVNs on γc-cytokine-induced cell proliferation in transfected Ba/F cells. (L) BaF-B03_IL2Rβ, (M) Ba/F3_IL4Rα, and (N) Ba/F3_IL7Rα cells were incubated with IL-3 and IL-2, IL-4, or IL-7, respectively, in the presence of indicated IAVNs for 24 hours. Cell viability was measured by 3-(4,5-dimethyl-2-thiazolyl)-2,5-diphenyltetrazolium bromide (MTT) assay. Data in (A–K) are representative of at least three independent experiments. Data in (L–N) are presented as mean ± standard division of the mean (SDM).

### IL-2-induced functional signal complex assembly requires an acidic environment of intercellular vesicles

We believe that our data in Figure 1 represent a previously unknown immunosuppressive mechanism of CQ needing further investigation. Therefore, we next investigated how IAVNs exert inhibitory effects on γc-cytokine signaling. First, we conducted dose-escalation studies using TPA-MAT cells ^40^, an IL-2-responsive T cell line. We observed a dose-dependent inhibitory effect of IAVNs on IL-2-induced STAT5 phosphorylation (Fig. 2A–C, Supplementary Fig. 1J/K). Second, we examined whether IAVNs inhibit the phosphorylation of JAK1 and JAK3, two protein tyrosine kinases associated with the IL-2Rβ and IL-2Rγ, respectively. JAK1 and JAK3 phosphorylation was significantly attenuated by IAVNs (Fig. 2D). These inhibitory effects were associated with a lack of IL-2-induced phosphorylation of IL-2Rβ and IL-2Rγ receptor subunits ^41^. Attenuated activation of the JAK/STAT pathway by IAVNs may be due to the receptor downregulation or reduced IL-2 binding. For this, we next used NanoLuc-labeled IL-2 to monitor the IL-2/IL-2R binding ^42^. IAVNs showed negligible effects on cell-surface IL-2 binding (Fig. 2E). Intracellular IL-2, indicating IL-2-induced IL-2/IL-2R complex translocation into the endosomal compartments, was only modestly (up to 10% decrease by AC) decreased by the IAVNs (Fig. 2F). We performed an immunoprecipitation assay to clarify further whether IAVNs affected a functional IL-2R complex formation made of the four key components, IL-2Rβ/IL-2Rγ/JAK1/JAK3, and checked the four proteins by western blots. The addition of IL-2 enabled the complex multiprotein formation. However, the IL-2-induced interaction between the IL-2 receptor multimer and JAK1/3 was significantly impaired following IAVN treatment (Fig. 2G). These results suggest that IAVNs inhibit IL-2-induced signal transduction by interfering with recruiting JAK1 and JAK3 to the IL-2 signal complex.

**Figure 2.**
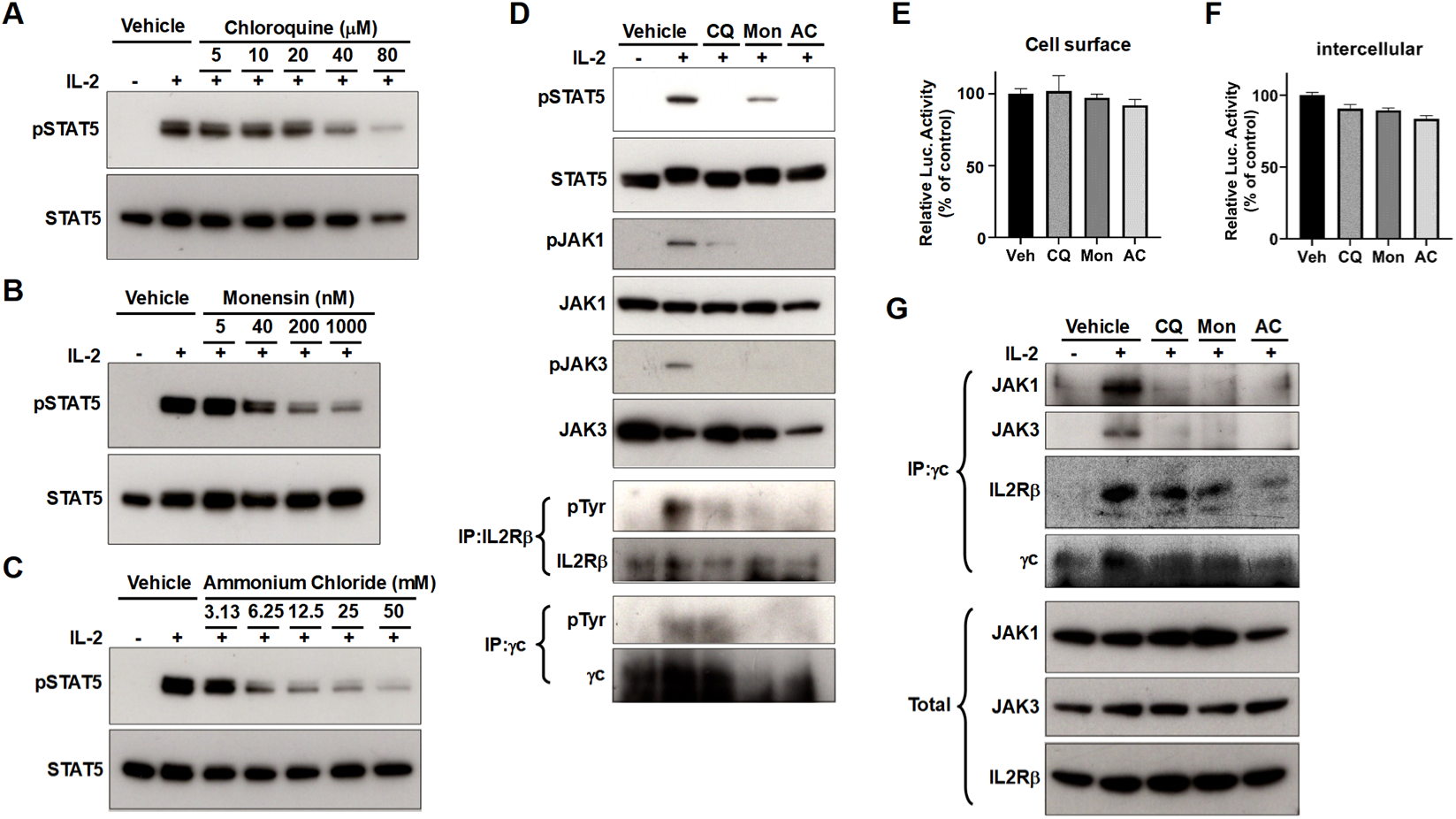
Intercellular acidic vesicle neutralizers (IAVNs) inhibit interleukin (IL)-2 induced recruitment of JAKs to the signal complex. (A–C) Dose-dependent suppressive effects of IAVNs on IL-2-induced STAT5 phosphorylation. TPA-MAT cells were pre-incubated with the indicated concentration of (A) chloroquine (CQ), (B) Monensin (Mon), or (C) ammonium chloride (AC) for one hour and stimulated with IL-2 for 10 minutes. (D) The effects of IAVNs on the phosphorylation of all components of the IL-2 signal complex. (E, F) The impact of IAVNs on (E) IL-2 binding on the cell surface and (F) endocytosis of IL-2. (G) The impact of IAVNs on IL-2-induced signal complex assembly. Cell lysates were immunoprecipitated with an anti-γc antibody and coprecipitates were analyzed by western blotting. Data in (A–D, G) are representative of at least three independent experiments. Data in (E, F) are presented as mean ± standard division of the mean (SDM).

### JAK3 targets the endosomal membrane in an acidic pH-dependent manner

The endosomal membrane facing the cytoplasm serves as a physical platform for signal complex assembly ^3–6^, with endosomal scaffolds and adaptor proteins contributing to signaling component recruitment from the cytoplasm ^43–46^. An acidic endosomal pH also affects membrane targeting of endosomal proteins via PtdIns3P ^11, 12^. To determine if JAKs can interact with the endosomal membrane in an acidic pH-dependent manner, nuclei, membrane, and cytosolic cell supernatant fractions were generated under neutral or acidic conditions. Interestingly, JAK3 was detected in the membrane fraction only under acidic conditions (Fig. 3A). This is similar to Hrs and EEA1, which are FYVE domain-containing endosomal proteins that interact with endosomal PtdIns3P in an acidic pH-dependent manner ^12^. In contrast, other JAKs were detected in the membrane fraction independently of pH. Furthermore, membrane-bound JAK3 under acidic conditions showed dissociation from the membrane following exposure to a neutral environment (Fig. 3B), again similar to Hrs and EEA1. We then separated the membrane fraction obtained under acidic conditions using density gradient centrifugation, finding that almost all JAK3 protein was detected in the Hrs and EEA1 concentrated fractions, and the fractionation pattern of JAK3 was similar to that of Hrs (Fig. 3C). These results consistently indicate that JAK3 binds the endosomal membrane in an acidic pH-dependent manner. Some evidence has demonstrated that γc is necessary for membrane binding of exogenously expressed JAK3 ^47–49^. We established a γc-knockout TPA-MAT cell line and confirmed the membrane binding capacity of JAK3 (Supplementary Fig. 2A/B). We detected JAK3 in the membrane fraction under acidic pH conditions in γc knockout TPA-MAT cells, as in wild type TPA-MAT cells (Supplementary Fig. 2C). Our results demonstrate that, unlike exogenously expressed JAK3, endogenous JAK3 interacts with the endosomal membrane in an acidic pH-dependent manner independently of γc.

**Figure 3.**
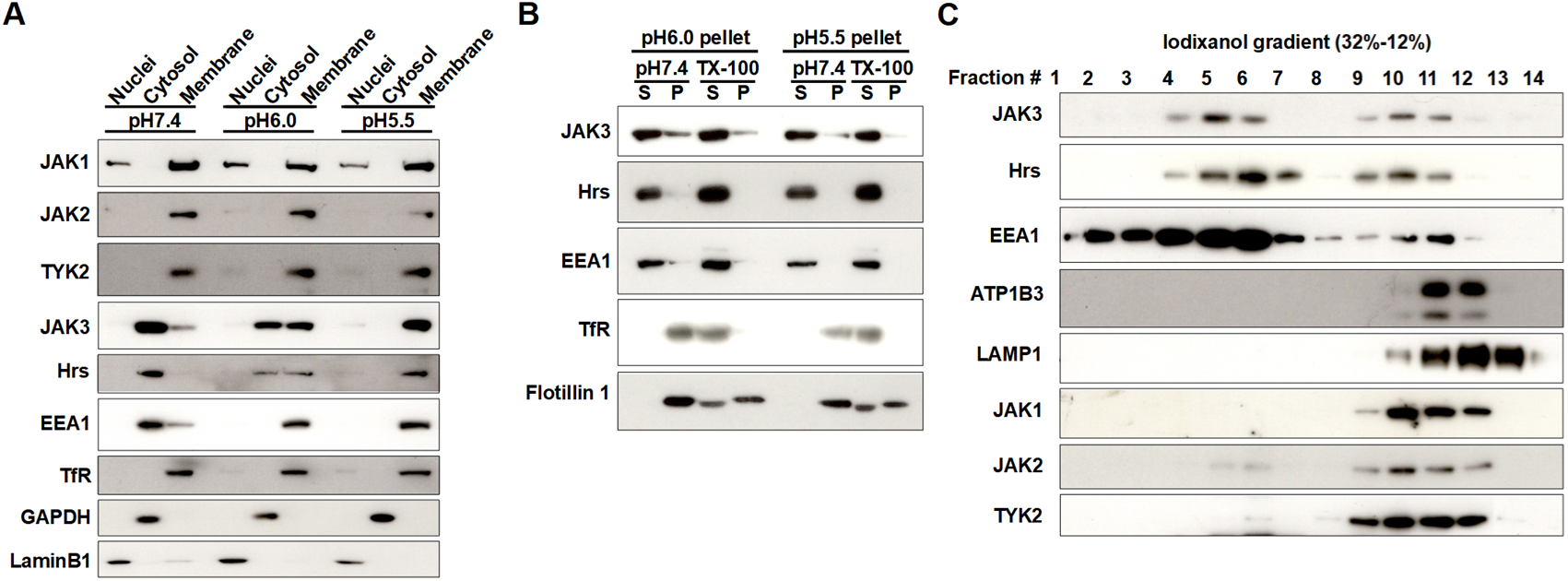
Acidic pH-dependent localization of JAK3 on the endosomal membrane. (A) Acidic pH-dependent membrane interaction of JAK3. Cells were homogenized at pH 7.4, pH 6.0, or pH 5.5 and fractionated into nuclei, membrane, and cytosol fractions by centrifugation. LaminB1, GAPDH, and Transferrin receptor (TfR) were used as nucleic, cytosolic, and membrane fraction markers, respectively. (B) JAK3 readily dissociates from the membrane fractions by exposure to a neutral environment. Membrane fractions at pH 6.0 and 5.5 were resuspended in homogenization buffer at pH 7.4 with or without of 1% TritonX-100, followed by re-fractionation into supernatant (S) and membrane pellet (M) by 100,000×g ultra-centrifugation. TfR and Flotillin1 indicate detergent soluble and insoluble membrane protein markers, respectively. (C) JAK3 interacts with endosomal membranes under an acidic pH. Membrane fractions at pH 6.0 were re-fractionated by density-gradient centrifugation. Hrs and EEA1, ATP1B3, and LAMP1 indicate endosome, plasma membrane, and lysosome fractions, respectively. All data are representative of at least three independent experiments.

### Recruitment of JAK1 and JAK3 to the IL-2 signal complex occurs after endocytosis

After IL-2 binding at the cell surface, IL-2 receptor multimers are endocytosed and sorted into endosomes. Our results suggest that IL-2-induced signal complex formation possibly occurs on endosomes. We therefore assessed the effect of inhibiting endocytosis. First, we assessed STAT5 phosphorylation. Dynasore, IPA3 and EHT1864, three previously verified IL-2 receptor endocytosis inhibitors^49–51^, consistently impaired IL-2-induced STAT5 phosphorylation (Supplementary Fig. 3A). We next examined the phosphorylation of IL-2 signaling complex proteins. Like STAT5, phosphorylation of JAK1 and JAK3 was attenuated by Dynasore treatment (Supplementary Fig. 3B). Accordingly, IL-2Rβ-and IL-2Rγ-phosphorylation was also severely reduced. Furthermore, like IAVNs, IL-2-induced functional signal complex formation was disrupted by Dynasore treatment (Supplementary Fig. 3C). These results suggests that the IL-2-induced endocytosis is required for the functional IL-2 signal complex formation.

### JAK3 is necessary for recruitment of JAK1 to the IL-2 signal complex

Like JAK3, IL-2-induced JAK1 incorporation into the IL-2 signal complex was disrupted by both IAVNs and endocytosis inhibitor treatment (Fig. 2G, Supplementary Fig. 3C), although JAK1 can bind the plasma membrane and endosomal membrane independently of acidic pH (Fig. 3A/C). These results imply that JAK1 recruitment to the IL-2 signal complex first requires JAK3 incorporation. We therefore performed respective knockdowns of JAK1 and JAK3. JAK1-knockdown severely reduced STAT5-phosphorylation and JAK3, IL-2Rβ, and γc (Supplementary Fig. 4A). JAK3-depletion almost totally diminished phosphorylation of all IL-2 signaling pathway components, STAT5, JAK1, IL-2Rβ, and γc. Nex, we performed co-IP experiments with γc. JAK1 knockdown had little if any effect on IL-2-induced IL-2Rβ/γ complex formation and JAK3 recruitment (Supplementary Fig. 4B). On the contrary, JAK3-depletion interfered with the IL-2-induced interaction between IL-2 receptor multimers and JAK1 (Supplementary Fig. 4C). These results suggest JAK3-incorporation to the IL-2 signal complex is required for that secondary recruitment of JAK1 to the IL-2 signal complex (Supplementary Fig. 4D).

**Figure 4.**
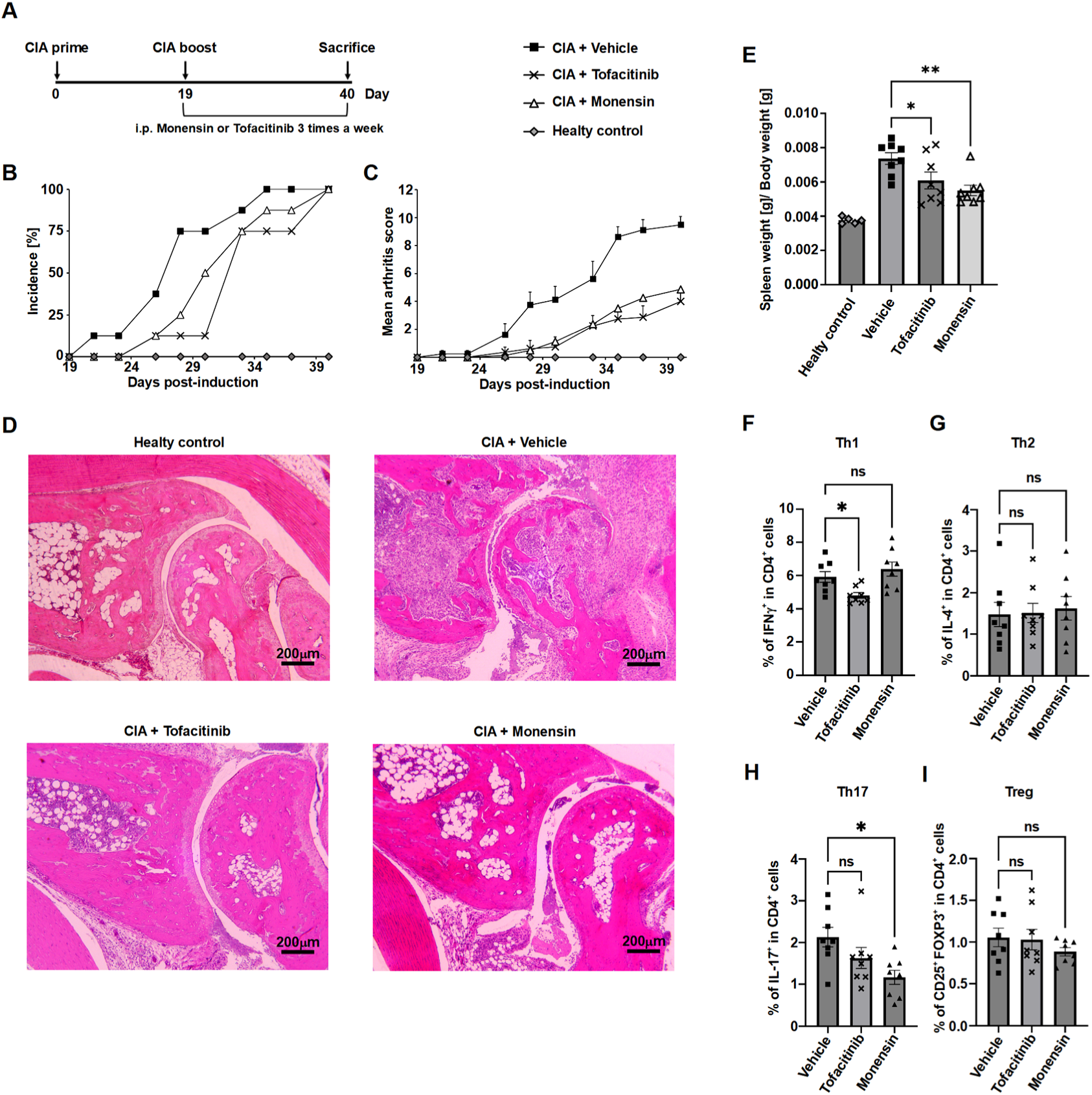
Anti-inflammatory effects of Monensin in a collagen-induced arthritis mouse model. (A) The study design for induction of arthritis in a mouse model. Immunized mice (n=24) were divided into three groups (n=8) and received vehicle, 25 mg/kg Tofacitinib, or 1 mg/kg Monensin thrice weekly. Unimmunized mice (n=5) were used as a healthy control group. (B) The incidence of arthritis (percentage of immunized mice). (C) The mean arthritic score for each group. (D) The representative histology of the affected joints. Hind paw sections were stained with hematoxylin and eosin. (E) The mean spleen weight (g) per body weight (g) for each group at day 40. (F–I) The mean percentage of (F) interferon (IFN) γ−positive cells, (G) interleukin (IL)-4-positive cells, (H) IL-17-positive cells, and (I) CD25 high and FOXP3-positive cells in CD4-positive splenocytes for each group. Data in (C, E– I) are presented as mean ± standard error of the mean (SEM). Statistical significance was determined by one-way ANOVA with Dunnett’s multiple comparison test (\**P*<0.05, \*\**P*<0.01).

### Monensin exerts a therapeutic effect on collagen-induced arthritis in mice

Our observation that carrier ionophores Monensin, Nigericin, and Salinomycin inhibit γc-cytokine-induced signal transduction prompted us to confirm their anti-inflammatory effects in an in vivo autoimmune model. We asked whether Monensin has any therapeutic potential in collagen-induced arthritis (CIA) mouse model (Fig. 4A). For positive control, we chose Tofacitinib, a JAK1 and JAK3 selective inhibitor known to be efficacious in Rheumatoid Arthritis (RA) ^52^. Tofacitinib administration retarded RA-onset and reduced the arthritis score (Fig. 4B, C). Treatment with Monensin significantly delayed the development of arthritis and reduced the disease severity. We next performed a pathological assessment of ankle joints (Fig. 4D). Similarly to Tofacitinib, Monensin treatment significantly suppressed inflammatory cell infiltration in the joint cavity, cartilage damage and bone degradation. It is of note that spleen enlargement, a hallmark often associated with an RA-related Felty syndrome symptom, was similarly suppressed by the two drugs (Fig. 4E). We further investigated whether Monensin possesses any effect on immune cell balance on CD4^+^ T cell populations. The mouse CIA model is one of the best-characterized animal models of RA, where Th1 and Th17 cells are the critical pathogenic T cell subsets ^53–55^. To this end, we examined Th1 and Th17 cells using splenocytes. Tofacitinib treatment significantly decreased the IFNγ-positive T-helper (Th) 1 cell population, but IL-17-positive Th17 cell reduction was only marginal (Fig. 4F/H). In contrast, Monensin treatment significantly decreased the population of Th17 cells, while it did not affect the population of Th1 cells. Neither of the two agents affected the population of Th2 cells and regulatory T (Treg) cells (Fig. 4G/I). Collectively, our results demonstrate that Monensin can modulate CIA inflammation by suppressing Th17 cells.

## Discussion

Small molecules that inhibit JAKs have gained traction as efficacious options for the treatment of autoimmune diseases. JAK inhibitors have excellent anti-inflammatory properties, so many have therefore been developed and are being used clinically to treat autoimmune diseases, allergic diseases, and COVID-19 infections ^24, 25, 56, 57^. JAK inhibitors interfere with the ATP-binding pocket of JAKs, which is essential for the catalytic activity ^24, 25^. Selective JAK3 inhibitors (JAK3i) have recently gained attention. Because JAK3 only regulates γc-cytokine signaling, it has become a potentially ideal target from both an efficacy and safety perspective ^26, 27^. Several JAK3i, including Ritlecitinib, have been developed, and it has been confirmed that they have sufficient inhibitory effects on γc-cytokine signaling to elicit anti-inflammatory activity ^58–61^. However, it has been reported that JAK1, not JAK3, phosphorylates STAT5 in IL-2-induced signal transduction ^62, 63^. Furthermore, it has been demonstrated that JAK3 catalytic activity is not necessary for IL-2-induced STAT5 phosphorylation, although a decreased intensity of STAT5 phosphorylation is associated with the loss of JAK3 activity ^63^. This evidence suggests that JAK3 catalytic activity is possibly not the best target for suppressing γc-cytokine signaling. In this study, we demonstrated a spatial regulation mechanism for γc-cytokine-induced signal transduction, which is a potential new target of anti-inflammatory drugs. We found that the endosomal localization of JAK3 is necessary for the recruitment of not only JAK3, but also JAK1 to the IL-2 signal complex, and is essential for γc-cytokine signaling. We propose that endosome neutralization is one way to interfere with this mechanism.

The subcellular localization of JAKs has previously been investigated using exogenous gene expression ^47, 64^. However, the subcellular localization of exogenously expressed proteins does not necessarily reflect the localization of endogenous proteins. In the present study, we performed cell fractionation using density gradient centrifugation to observe the subcellular localization of endogenous JAKs. It has been reported that overexpressed JAK1 and JAK2 are localized on the plasma membrane ^64^, which is consistent with our results for endogenous JAK1 and JAK2. However, overexpressed JAK3 reportedly diffuses in the cytoplasm ^47–49^. This differs from our finding that endogenous JAK3 is present in the endosomal fractions in an acidic pH-dependent manner. This difference is likely because the endosomal localization of JAK3 is restricted by the abundance of endosomal scaffold and adapter proteins that interact with JAK3. The TGFβ/Smad pathway, a well-established model of endosome-specific signal activation, reinforces our hypothesis. Smad Anchor for Receptor Activation (SARA), a FYVE domain-containing protein that localizes to the endosomal membrane in an acidic pH-dependent manner, directly interacts with Smad2/3 at endosomal membranes. Once activated, the TGFβ receptor is transported into the endosome, where it can phosphorylate Smad2/3 efficiently through its interaction with SARA ^43, 44, 65, 66^. As we observed with JAK3, exogenously expressed Smad2/3 diffuses in the cytoplasm because of insufficient levels of SARA, although endogenous SMAD2/3 was observed on the endosome ^43, 44^. We believe that JAK3 endosomal localization is mediated by adaptor or scaffold proteins that interact with the endosomal membrane in an acidic pH-dependent manner, such as FYVE domain-containing proteins ^9, 11, 12^. The specific identification of such molecules could contribute to understanding of γc-cytokine-mediated signaling mechanisms and help discover new molecular targets for anti-inflammatory drugs.

When exogenous JAK3 is co-overexpressed with γc, JAK3 can co-localize with γc at the plasma membrane via a constitutive interaction ^41, 47^. In contrast, interactions between endogenous JAK3 and γc were scarcely observed without ligand stimulation ^33, 67^. Using γc knockout cells, we confirmed that γc was unnecessary for membrane binding of endogenous JAK3. Furthermore, we verified that low levels of endogenous JAK3 were detected in the plasma membrane fractions. The significance of the constitutive interaction between JAK3 and γc at the plasma membrane remains unclear.

Our results demonstrate that carrier ionophores that act as proton/cation exchangers, including Monensin, can suppress γc-cytokine-induced cell proliferation with high sensitivity and selectivity compared with CQ. Monensin repetitively carries protons out of acidic vesicles until it is metabolized ^38^, while protonated CQ is trapped and accumulates in the acidic vesicles ^14, 15^. Although our validation is limited, such a difference in mechanism of action may significantly affect the efficacy and selectivity of these molecules. Carrier ionophores have not been used in humans but are widely used as livestock feed additives and their safety has been established in several animals ^68^. This suggests that carrier ionophores could be anti-inflammatory drugs that are superior to CQ. Therefore, we verified the anti-inflammatory effects of Monensin using CIA mice. Monensin could suppress CIA comparably to Tofacitinib, which has high inhibitory activity against JAK1 and JAK3 ^52^. However, we observed significantly different effects on T cell subsets between them. Monensin exhibited a more potent suppressive effect on Th17 cells than Tofacitinib. Unlike Tofacitinib, Monensin does not inhibit IFN signaling, which suppresses Th17 cell differentiation ^69, 70^, possibly explaining our results. Monensin did not affect Th1 cells, similar to CQ treatment as reported by others ^71^. Notably, it has been reported that JAK3i can suppress Th1 cells similarly to Tofacitinib ^59–61^. We believe that the different impacts on Th1 cells between Monensin and JAK3i are caused by their off-target effects. JAK3i were designed to target Cys 909 in the ATP binding pocket of JAK3, which is structurally different from other JAKs. In contrast, JAK3i may possibly inhibit kinases, including TEC-family kinases, which have a Cys residue at the equivalent position of Cys 909 in JAK3 ^58^. Monensin-induced endosomal neutralization may influence the signaling of some cytokines that can interfere with inflammatory responses. Indeed, we observed that IAVNs could consistently inhibit GM-CSF signaling. In addition, endosomal neutralization might augment signal activation ^72, 73^. The reason why Monensin, unlike JAK3i, does not affect Th1 cells is unclear, while this indicates that Monensin could potentially be an anti-inflammatory drug like CQ without an associated increased risk of infection. ^74^

Overall, we demonstrated the effects of IAVNs on γc-cytokine-induced signal transduction via endosomes. The neutralization of acidic vesicles also affects lysosomal functions, which support critical inflammatory events, including the processing of antigens for presentation on MHC molecules ^19, 20^, activation of some TLRs by proteolytic cleavage, and autophagy-regulated inflammatory cytokine secretion ^14, 21, 22, 75^. Although the anti-inflammatory effects of IAVNs are attributed to these combined effects, we found that the inhibition of γc-cytokine-induced signal transduction is undoubtedly an important mechanism of action. Limitations of research While various immune cells are involved in autoimmune inflammation, our analysis is limited to lymphocytes. Analysis using PBMCs revealed that GM-CSF-induced STAT activation, which mainly arises in monocytic lineage cells, was inhibited by IAVNs treatment despite JAK3 not being required for GM-CSF-induced signal transduction. Further studies are needed to clarify the whole role of the endosomal environment in signal transduction. Additionally, the scaffolding or adapter proteins that mediate JAK3-targeting to endosomes should be identified in future work. This will further clarify the molecular mechanism of JAK3-mediated signal transduction via endosomes.

## Acknowledgments

This work was supported by the JSPS KAKENHI grant (21K08474 and 20K08203) and AMED grant (JP22ym0126802 and 21cm0106387h0001). We thank the Research Center for Supports to Advanced Science, Shinshu University for use of facilities and M. Amano for technical support in preparing texture sections. We thank J. Iacona, Ph.D., from Edanz (https://jp.edanz.com/ac) for editing a draft of this manuscript.

## Author contributions

Y.A. designed the research; Y.A., N.T., T.T., and K.K. wrote the paper; Y.A. performed the in vitro experiments; Y.A. and K.K. performed the in vivo experiments.

## Methods

Experimental model and subject details

### Human samples

The study was approved by the ethics committee of Shinshu University School of Medicine (approval number: 5625), and written informed consent was obtained from all subjects. PBMCs were obtained from heparinized blood by centrifugation (800×g for 20 min) using PBMC spin medium (PluriSelect, Cat. # 60-00092-12).

### Cell culture

The mouse IL-3-dependent pro-B cell lines, Ba/F3, and BaF-B03_IL2Rβ ^37^, were maintained in RPMI-1640 (Fujifilm, Cat. # 189-02025) supplemented with 10% fetal bovine serum (FBS), 5 μM mercaptoethanol, 1× antibiotics (Fujifilm, Cat. # 168-23191), and 10 ng/mL recombinant mouse IL-3 (Peprotech, Cat. # 213-13). TPA-MAT cells ^40^ were maintained in RPMI-1640 supplemented with 10% FBS, 1× antibiotics, and 100 ng/mL phorbol 12, 13-dibutyrate (PDBu) (Fujifilm, Cat. # 198-09752). HEK293T cells were grown in high glucose DMEM (Fujifilm, Cat. # 043-30085) supplemented with 10% FBS and 1× antibiotics. All cells were cultured at 37°C and 5% CO_2_.

### Mice

DBA/1JJmsSlc mice (seven weeks old) were purchased from Japan SLC. All animal procedures were performed according to the guidelines of the Animal Care and Use Committee of Shinshu University following a protocol approved by the committee (approval number: 021025). All mice were kept under specific pathogen-free conditions.

## Method Details

### Transfection and infection of cells

The coding sequences of human IL4Rα and human IL7Rα were each inserted into the pCDH-CMV-MCS-EF1-Puro vector (System Biosciences, Cat. # CD510B-1) between the EcoRI and NotI restriction sites. Short hairpin RNA (shRNA) sequences targeting JAK1 and JAK3 were inserted into the pRSI12-U6-sh-HTS4-UbiC-TagRFP-2A-puro vector (Cellecta) at the BbsI restriction site. The shRNA sequences were acquired from DECIPHER Lentiviral shRNA Library (Cellecta). The guide RNA sequence targeting exon 1 of γc was inserted into the lentiCRISPR v2 vector (addgene Cat. # 52961) at the BsmBI restriction site. Lentivirus production utilized HEK293T as the producer cell line. HEK293T cells (6 cm dish, 90% confluent, approximately 3x10^6^ cells) were co-transfected with 1 μg of each lentiviral construct (pCDH-puro-hIL4Rα, pCDH-puro-hIL7Rα, pRSI12-hJAK1KD, pRSI12-hJAK3KD, pRSI12-U6-sh-HTS4-UbiC-TagRFP-2A-puro, or lentiCRISPRv2-γc), 1 μg of pMDLg/pRRE, 1 μg of pRSV-Rev, and 300 ng of pMD2.G (addgene, Cat. # 12251, 12253, 12259) using 20 μg of PEI MAX® (Polysciences, Cat. # 49553-93-7) in 3 mL of culture media. After 12 hours, transfection media were replaced with 2 mL of pre-warmed fresh media. After 24 hours, cell culture media containing lentiviral particles were collected and filtered with Millex®-HV 0.45 μm (Merck, Cat. # SLHVR33RB). Ba/F3 and TPA-MAT cells were cultured with lentiviral particle-containing media in the presence of 8 μg/mL Polybrene (Nacalai tesque, Cat. # 12996-81). After 8 hours, cells were washed with PBS and cultured with the culture media described above. After 24 hours, infected cells were selected with 2 μg/mL puromycin (Sigma-Aldrich, Cat. # P8833) for 2 days. Single cell clones of γc knockout TPA-MAT cells were obtained by limiting dilution.

### Cytokine-induced STAT phosphorylation in PBMCs

Freshly isolated PBMCs were pre-incubated with 80 μM CQ, 1 μM Monensin, 50 mM ammonium chloride, or Vehicle (0.02% methanol) in RPMI-1640 supplemented with 10% FBS and 5 μM mercaptoethanol for 1 hour (1x10^6^ cells/mL). Cells were then stimulated with 15 ng/mL IL-2 (Shionogi), 10 ng/mL IL-7 (Peprotech, Cat. # 200-007), 10 ng/mL IL-15 (Biolegend, Cat. # 570302), 1 ng/mL GM-CSF (Peprotech, Cat. # 300-003), 10 ng/mL IL-6 (Peprotech, Cat. # 200-006), 10 ng/mL IL-10 (Peprotech, Cat. # 200-010), 2 ng/mL IFNα (Biolegend, Cat. # 592702), or 10 ng/mL IFNγ (Peprotech, Cat. # 300-02) for 30 minutes. Cells were collected by centrifugation at 400×g for 1 minute, then lysed for 20 minutes on ice in cell lysis buffer containing 1% NP-40, 20 mM Tris-HCl (pH7.4), 150 mM NaCl, 1 mM EDTA, 2.5 mM sodium pyrophosphate, 1 mM β-glycerophosphate, 1 mM sodium orthovanadate, and 1% protease inhibitor cocktail (Nacalai tesque, Cat. # 25955-11). Cellular debris was pelleted by centrifugation at 20,000×g at 4°C for 20 minutes. Cell extracts were analyzed by western blotting, as described below.

### IAVN treatment of cell lines

Transfected TPA-MAT and Ba/F cells were washed with PBS twice, then pre-cultured in RPMI-1640 supplemented with 10% FBS for 4 hours. In Figure 1I/J, Figure 2D/G, and Supplementary Figure 1D– F, cells were pre-incubated with 80 μM CQ, 1 μM Monensin, 50 mM ammonium chloride, 1 μM Nigericin, 1 μM Salinomycin, or vehicle (0.02% methanol) in RPMI-1640 supplemented with 10% FBS and 5 μM mercaptoethanol for 1 hour (1x10^6^ cells/mL). In Figure 2A–C and Supplementary Figure 1J/K, cells were pre-incubated with IAVNs at the indicated concentrations in RPMI-1640 supplemented with 10% FBS and 5 μM mercaptoethanol for 1 hour (1x10^6^ cells/mL). After pre-incubation, cells were stimulated with 15 ng/mL human IL-2, 1 ng/mL human IL-4, 20 ng/mL human IL-7, or 2 ng/mL mouse IL-3 for 10 minutes. Cells were collected by centrifugation at 400×g for 1 minute, then lysed with cell lysis buffer (1x10^7^ cells/mL) on ice for 20 minutes. Cellular debris was pelleted by centrifugation at 20,000×g at 4°C for 20 minutes. Cell lysates were analyzed by western blotting or used in immunoprecipitation assays, as described below.

### Cell proliferation assay

The effect of IAVNs on cytokine-induced cell proliferation was evaluated using tetrazolium salt WST-1 (Roche, Cat. # 5015944001). Transfected Ba/F cells were washed with PBS twice, then pre-cultured in RPMI-1640 supplemented with 10% FBS for 4 hours. Then, cells were seeded into 96-well culture plates at 20,000 cells/well in 100 μL RPMI-1640 supplemented with 10% FBS, 5 μM mercaptoethanol, and the indicated concentration of IAVNs (CQ: 5–160 μM, Monensin: 1.6 nM–5 μM, Ammonium Chloride: 1.56–50 mM, Nigericin: 100 pM–10 μM or Salinomycin: 1.6 nM–5 μM). Cells were incubated in the presence of 15 ng/mL human IL-2, 1 ng/mL human IL-4, 20 ng/mL human IL-7, or 2 ng/mL mouse IL-3 for 24 hours. All plates included blank wells without cells for correction of measurement values. Each condition included four replicates. To measure cell proliferation levels, 10 μL of WST-1 was added to each well and mixed gently, then incubated for 1 hour at 37°C. The absorbance was measured at 440 nm (measurement wavelength) and 650 nm (control wavelength) using a SpectraMax iD5 instrument (Molecular devices). Correction of each sample absorbance value (440 nm) was performed by subtracting the absorbance values of both the respective control wavelength (650 nm) and average of the blank wells (440 nm - 650 nm). The relative inhibition rate was calculated by making comparisons with the average absorbance values of the non-treated wells. The EC50 value was determined by fitting a four-parameter log-logistic model.

### IL-2 binding and endocytosis assay

To assess the effects of IAVNs on IL-2 binding to the cell surface receptor complex and endocytosis, we used NanoLuc-IL-2, which contains G4S flexible linker and NanoLuc® luciferase (Promega) at the C-terminus of IL-2, as previously reported ^42^. TPA-MAT cells were washed with PBS twice, then pre-cultured in RPMI-1640 supplemented with 10% FBS for 4 hours. Then, cells were pre-incubated with 80 μM CQ, 1 μM Monensin, 50 mM ammonium chloride, or vehicle (0.02% methanol) in RPMI-1640 supplemented with 10% FBS for 1 hour (1x10^6^ cells/mL). To analyze IL-2 binding to the cell surface receptor, pre-incubated cells (1x10^4^) were collected and re-suspended with 50 μL of pre-chilled media containing the same IAVNs that were used in pre-incubation and stored on ice for 10 minutes. The cells were incubated with 100 pM NanoLuc-IL-2 on ice for 30 minutes, then washed with ice-cold PBS and lysed with passive lysis buffer (Promega, Cat. # E1941). To analyze endocytosis of IL-2, pre-incubated cells (1x10^4^ cells/50 μL) were incubated with 100 pM NanoLuc-IL-2 at 37°C for 10 minutes. Subsequently, cells were washed with PBS and lysed with passive lysis buffer.

The NanoLuc luciferase assay was performed using the Nano-Glo^®^ Luciferase Assay System (Promega, Cat. # N1110) and SpectraMax iD5 following the manufacturer’s instructions.

### Cell fractionation

Cells (1x10^7^) were washed with PBS and re-suspended in 1 mL of ice-cold homogenization buffer (20 mM HEPES (Ph7.4), MES (pH 6.0) or MES (pH 5.5), 250 mM sucrose, 100 μM EDTA, 2%Protease inhibitor cocktail, and 20 μM E-64-d (Peptide institute, Cat. # 4321-v)). Homogenization was performed using a 26 Gauge needle and 1 mL syringe. The cell suspension was passed through the syringe repeatedly until approximately 80% of cells were disrupted. Homogenates were centrifuged two times at 400×g for 10 minutes at 4°C to eliminate debilis and undisrupted cells. Homogenates were re-centrifuged at 1,000×g for 30 minutes at 4°C and pellets were collected and labeled as the nuclei fraction. Then, the supernatants were ultra-centrifuged with an angle rotor at 100,000×g for 30 minutes at 4°C to separate the cytosolic fraction (supernatant) and membrane fraction (pellet). The nuclei and membrane pellets were dissolved in PBS containing 1% SDS in an equal volume of cytosolic fraction. Density gradient fractionation was performed using OptiPrep™, which contains 60% iodixanol. To generate an iodixanol concentration gradient, we prepared a 32%–12% iodixanol solution, which contains 20 mM MES (pH 6.0), 100 μM EDTA, 2% Protease inhibitor cocktail, and 20 μM E-64-d in 2.5% increments by diluting with homogenization buffer (pH 6.0). Each 1.6 mL of iodixanol solutions were layered in descending order of iodixanol concentration in centrifuge tubes (Beckman Coulter, Cat. # 344059) and left to stand for 24 hours at 4°C to generate a continuous density gradient. The membrane pellet obtained under the pH 6.0 condition described above was suspended in 200 μL of 12% iodixanol solution completely and laid on the iodixanol gradient, then centrifuged at 100,000×g for 16 hours at 4°C using a SW-41-Ti swing rotor (Beckman Coulter). Then, separated samples were collected at the bottom of centrifuge tubes (600 μL) for each fraction. To analyze pH-dependent protein dissociation, membrane fractions (pellets) obtained from pH 6.0 and pH 5.5 samples were suspended in homogenization buffer (pH 7.4) with or without of 1% Triton-X by pipetting. After incubation for 10 minutes on ice, samples were centrifuged at 100,000×g for 30 minutes at 4°C to separate the supernatant containing dissociated membrane proteins under neutral pH and the membrane pellet. The pellet was dissolved in PBS containing 1% SDS in an equal volume of supernatant. For western blot analysis, all protein samples obtained by cell fractionation were purified by methanol-chloroform precipitation. Briefly, one volume of sample was mixed with four volumes of methanol and vortexed well. The samples were mixed with one volume of chloroform and vortexed well again, then subsequently mixed with three volumes of water, vortexed for one minute, and centrifuged at 20,000×g for 10 minutes at room temperature. After removing of the upper layer, the middle and lower layers were rinsed with three volumes of methanol and dissociated completely. The pellet was dissolved with SDS-PAGE sample buffer (50 mM Tris-HCl (pH 6.7), 2% SDS, 10% glycerol, 2% 2-mercaptoethanol, and 0.02% bromophenol blue) and boiled for five minutes.

### Inhibition of endocytosis

TPA-MAT cells (5x10^5^) were washed with PBS three times and seeded into 6 cm culture dishes in 3 mL of pre-warmed serum-free RPMI-1640. After a five-minute incubation, the media were replaced with 3 mL of RPMI-1640 containing 5 μM mercaptoethanol and 80 μM Dynasore (Cayman Chemical, Cat. # 14062), 20 μM IPA3 (Cayman Chemical, Cat. # 14759), 10 μM EHT1864 (Cayman Chemical, Cat. # 17258), or 0.1% DMSO and pre-incubated for 30 minutes at 37°C. Afterwards, 50 ng/mL IL-2 was added to the dish and incubated for 10 minutes at 37°C. Then, media were discarded completely and cells were lysed with cell lysis buffer for 20 minutes on ice. Cellular debris was pelleted by centrifugation at 20,000×g at 4°C for 20 minutes. Cell lysates were analyzed by western blotting, as described below. To prepare the samples for immunoprecipitation assays, we seeded 1x10^7^ cells into 15 cm culture dishes in 20 mL of medium.

### Immunoprecipitation

The cell lysates (1x10^7^cells/mL) described above were filtered with Millex®-HV 0.45 μm before immunoprecipitation. To detect the tyrosine phosphorylation levels of IL2Rβ and γc, 200 μL and 1 mL of cell lysate, respectively, were used. Lysates were rotated with 3 μg of mouse anti-human IL2Rβ antibody (TU11) or 3 μg of rat anti-human γc antibody (TUGh4) and 10 μL of Dynabeads Protein G (Thermo Fisher, Cat. # DB10004) at 4°C for 3 hours. After that, Dynabeads were washed with 1mL of Lysis buffer three times and eluted by boiling for five minutes in SDS-PAGE sample buffer. To observe IL-2 signal complex formation, 2 mL of cell lysates were rotated with 5 μg of TUGh4 and 10 μL of Dynabeads Protein G at 4°C for 16 hours. Then, Dynabeads were washed with cold wash buffer (20 mM Tris-HCl (pH 7.4), 500 mM NaCl, 1mM EDTA, and 0.02% Tween-20) four times, followed by rinsed with PBS containing 0.02% Tween-20. Immunoprecipitates were eluted by boiling for five minutes in SDS-PAGE sample buffer. Supernatants were analyzed by western blotting, as described below.

### Western blotting

The cell lysates described above were boiled for five minutes in SDS-PAGE sample buffer, then separated using a 4-15% Mini-PROTEIN® Precast Gel (Bio-Rad, Cat. # 4561086). Proteins were transferred to Immobilon® Transfer Membrane (Millipore, Cat. # IPVH00010), and the membrane was blocked with 5% skim milk in TBST (20 mM Tris-HCl (pH 7.6), 150 mM NaCl, and 0.1% Tween-20) for 1 hour at room temperature. Incubation with primary antibodies, except anti-Phospho-Tyrosine (P-Tyr-1000, Santa Cruz Biotechnology, Cat. # 8954) and anti-Phospho-JAK3 (Cell Signaling Technology, Cat. # 5031), was performed in 5% skim milk in TBST with 0.5 μg/mL of each primary antibody at 4°C for 12 hours. Incubation with anti-Phospho-Tyrosine was performed in Blocking One-P (Nacalai tesque, Cat. # 05999-84) with 0.5 μg/mL anti-Phospho-Tyrosine antibody at 4°C for 12 hours. Incubation with anti-Phospho-JAK3 was performed in 1% BSA in TBST with 0.5 μg/mL anti-Phospho-JAK3 antibody at 4°C for 16 hours. Membranes other than those incubated with the anti-Phospho-Tyrosine antibody were incubated with horseradish peroxidase (HRP)-linked secondary antibodies in 5% skim milk in TBST at a 1:10,000 dilution for 1 hour at room temperature. Membranes incubated with the anti-Phospho-Tyrosine antibody were incubated with HRP-linked secondary antibody in Blocking one-P at a 1:100,000 dilution for 1 hour at room temperature. After thorough washing with TBST, signal was visualized using the Immobilon™ Western Chemiluminescent HRP Substrate (Millipore, Cat. # WBKLS0500) and Hyperfilm™ ECL™ (Cytiva, Cat. # 28906840). To reprobe the membrane, HRP was first deactivated by incubation with 34.5% hydrogen peroxide for 30 minutes at room temperature. Antibody stripping was performed using WB Stripping Solution (Nacalai tesque, Cat. # 05364-55) following the manufacturer’s instructions. The following antibodies were used for western blotting: anti-phospho-STAT5 (Tyr694), anti-STAT5, anti-phospho-STAT3 (Tyr705), anti-STAT3, anti-phospho-STAT1, anti-STAT1α p91, anti-phospho-STAT6 (Tyr641), anti-STAT6, anti-phospho-JAK1 (Tyr1034/1035), anti-JAK1, anti-phospho-JAK3 (Tyr980/981), anti-JAK3, anti-IL2Rβ, anti-IL2Rγ, anti-JAK2, anti-TYK2, anti-Hrs, anti-EEA1, anti-CD71 (TfR), anti-GAPDH, anti-Lamin B1, anti-Flotillin 1, anti-phospho-Tyrosine (P-Tyr-1000), anti-ATP1b3, anti-LAMP1, and anti-HSP60.

### Induction, treatment, and assessment of CIA

Emulsion was prepared by mixing bovine type II collagen (Chondrex, Cat. # 20022) and an equal volume of complete Freund’s adjuvant (Sigma-Aldrich, Cat. # F5881) in two-syringe method. After one week of habituation in the laboratory, DBA/1J mice (eight weeks old) were injected with 100 μL of the emulsion intradermally at the base of the tail. Day 0 was designated as the day of initial immunization. On day 19, mice were re-injected with 100 μL of emulsion consisting of bovine type II collagen and incomplete Freund’s adjuvant (Sigma-Aldrich, Cat. # F5506) intradermally at the lower back to boost the induction of CIA. On the same day (Day 19), drug administration and scoring the severity of paw inflammation were performed three times a week until the day of sacrifice (day 40). Each mouse was treated with 500 μL of 25 mg/kg Tofacitinib (Selleck, Cat. # S2789), 1 mg/kg Monensin, or saline intraperitoneally. The clinical arthritis scoring was performed in each limb on a scale of 0–4 and the total score of the limbs was evaluated. Scoring criteria were as follows: 0) No evidence of erythema and swelling, 1) Erythema and mild swelling confined to the tarsals or ankle joint, 2) Erythema and mild swelling extending from the ankle to the tarsals, 3) Erythema and moderate swelling extending from the ankle to the metatarsal joints, and 4) Erythema and severe swelling encompassing the ankle, foot, and digits, or ankylosis of the limb ^53^. Clinical evaluation was performed by two investigators together. Following sacrifice on day 40, the ankle joints and spleen were removed. Ankle joints were histologically examined, as described below. The spleen was weighed and analyzed with flow cytometry, as described below.

### Histological examination

The ankle joint was fixed with 4% paraformaldehyde for 24 hours, and the skin was stripped and subsequently re-fixed with 4% paraformaldehyde for 5 days. The ankle was degreased in 100% ethanol for 24 hours, then decalcified in 500 mM EDTA (pH 7.6) for 7 days. The ankle was washed with water and 100% ethanal, then embedded in a paraffin block. The 4 μm paraffin section was stained with hematoxylin and eosin.

### Flow cytometry

The protein expression levels of exogenous IL2Rβ, IL4Rα, and IL7Rα in transfected Ba/F cells were examined by flow cytometry. Cells (1x10^6^) were incubated with ice-cold FACS buffer (PBS containing 2% FBS and 1 mM EDTA) on ice for 15 minutes, then incubated with an anti-IL2Rβ antibody (TU11), anti-IL4Rα antibody (R&D Systems, Cat. # MAB230), or anti-IL7Ra antibody (BioLegend, Cat. # 351302) for 30 minutes on ice. After three washes with pre-chilled FACS buffer, the cells were incubated with a Goat anti-Mouse IgG (H+L) Superclonal™ Recombinant Secondary Antibody and Alexa Fluor™ 488 (Themo Fisher Scientific, Cat. # A28175) for 30 minutes on ice. After washing, the cells were fixed with 3% paraformaldehyde in PBS and stored at 4°C in the dark.

Splenocytes were removed from the spleen using a bended needle and filtered through a mesh strainer (70 μm) to generate single cell suspensions. Red blood cells were lysed with ACK lysis buffer (155 mM NH_4_Cl, 10 mM KHCO_3_, and 100 μM EDTA) (pH 7.3) on ice for 5 minutes, then washed twice with ice-cold PBS. For Treg assessment, splenocytes (2x10^6^) were first stained with 1:1,000 Zombie violet™ fixable viability kit (Biolegend, Cat. # 423113) in PBS at room temperature for 15 minutes. Then, cells were washed twice with pre-chilled FACS buffer and incubated with TruStain FcXTMPLUS anti-CD16/32 (BioLegend, Cat. # 156604) on ice for 15 minutes. The samples were stained for cell surface proteins (CD3, CD4, CD8, and CD25) using respective antibodies diluted in FACS buffer for 30 minutes on ice. The cells were then washed twice with FACS buffer and fixed and permeabilized using Foxp3/Transcription factor staining buffer kit (Themo Fisher Scientific, Cat. # 00-5523-00) following the manufacturer’s instructions. Foxp3 was stained for 30 minutes at room temperature. After washing, cells were resuspended with FACS buffer containing 1% paraformaldehyde and stored at 4°C in the dark. To detect cytokine production, splenocytes (2x10^6^) were incubated in RPMI-1640 containing 10% FBS, Cell activation cocktail (BioLegend, Cat. # 423302), and Protein transport inhibitor (BD Biosciences, Cat. # 51-2301KZ) for six hours at 37°C and 5% CO_2_. Cells were processed for Fc receptor blocking as described above. Then, staining for cell surface proteins (CD3, CD4 and CD8) was performed for 30 minutes on ice. Cells were washed twice with ice-cold FACS Buffer and fixed and permeabilized with Fixation buffer (BioLegend, Cat. # 420801) and Intercellular staining permeabilization wash buffer (BioLegend, Cat. # 421002) following the manufacturer’s instructions. The cells were then stained with anti-cytokine antibodies (IFNγ, IL-4, or IL-17) for 30 minutes at room temperature. After washing, cells were resuspended with FACS buffer containing 1% paraformaldehyde and stored at 4°C in the dark. All cells were analyzed with FACS Celesta™ and Flowjo software (BD Biosciences). The following antibodies were used for staining of splenocytes: anti-CD3-APC (17A2h), anti-CD4-APC-eFluor™780 (RM4-A5), anti-CD8-Super Bright™780 (53-6.7), anti-CD25-Alexa™488 (PC61.5), anti-FOXP3-PE-Cyanine7 (FJK-16s), anti-IFNγ-Brilliant Violet 605 (XMG1.2), anti-IL-4-PE-Cyanine7 (11B11), and anti-IL-17-Alexa488™ (TC11-18H10.1). All antibodies were used at a 2.5 μg/mL final concentration.

### Statistical analysis

All statistical analyses and graphing were performed using GraphPad Prism 9 (GraphPad). All statistical significance was determined by one-way ANOVA with Dunnett’s multiple comparison test (\**P*<0.05, \*\**P*<0.01, ns = not significant).

---

**Supplementary Figure 1.**
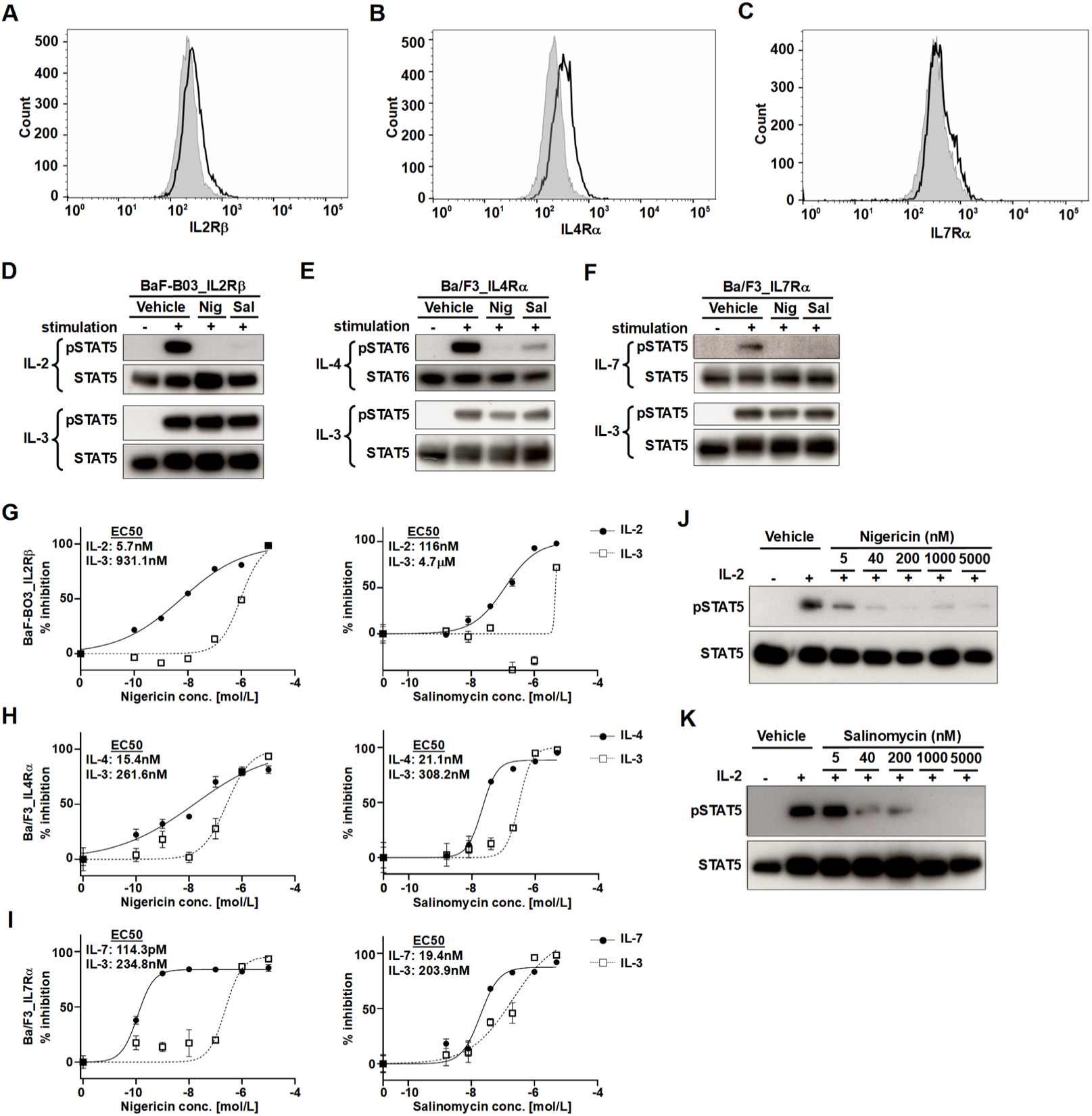
Carrier ionophores, Nigericin and Salinomycin, selectively interfere with γc-cytokine-induced signal transduction and cell proliferation. (A–C) The expression profiles of (A) IL-2Rβ, (B) IL-4Rα, and (C) IL-7Rα in BaF-BO3_IL2Rβ, Ba/F3_IL-4Rα, and Ba/F3_IL-7Rα cells, respectively, were assessed by flow cytometry. Filled histograms indicate non-transfected cells. (D–F) Selective suppression of carrier ionophores on γc-cytokine-induced STAT phosphorylation in transfected Ba/F cells. (D) BaF-B03_IL2Rβ, (E) Ba/F3_IL4Rα, and (F) Ba/F3_IL7Rα cells were pre-incubated with 1 μM Nigericin or 1 μM Salinomycin for one hour, then stimulated with interleukin (IL)-3 and IL-2, IL-4, or IL-7, respectively, for 10 minutes. Phosphorylation levels of STATs were analyzed by western blotting. (G–I) Selective inhibitory effects of carrier ionophores on γc-cytokine-induced cell proliferation in transfected Ba/F cells. (G) BaF-B03_IL2Rβ, (H) Ba/F3_IL4Rα, and (I) Ba/F3_IL7Rα cells were incubated in the presence of indicated carrier ionophores for 24 hours. Cell viability was measured by 3-(4,5-dimethylthiazol-2-thiazolyl)-2,5-diphenyltetrazolium bromide (MTT) assay. (J, K) Dose-dependent suppressive effects of carrier ionophores on IL-2-induced STAT5 phosphorylation. TPA-MAT cells were pre-incubated with indicated concentrations of (J) Nigericin and (K) Salinomycin for one hour and stimulated with IL-2 for 10 minutes. Phosphorylated STAT5 protein levels were assessed by western blotting. Data in (D–F, J, K) are representative of at least three independent experiments. Data in (G–I) are presented as mean ± standard division of mean (SDM).

**Supplementary Figure 2.**
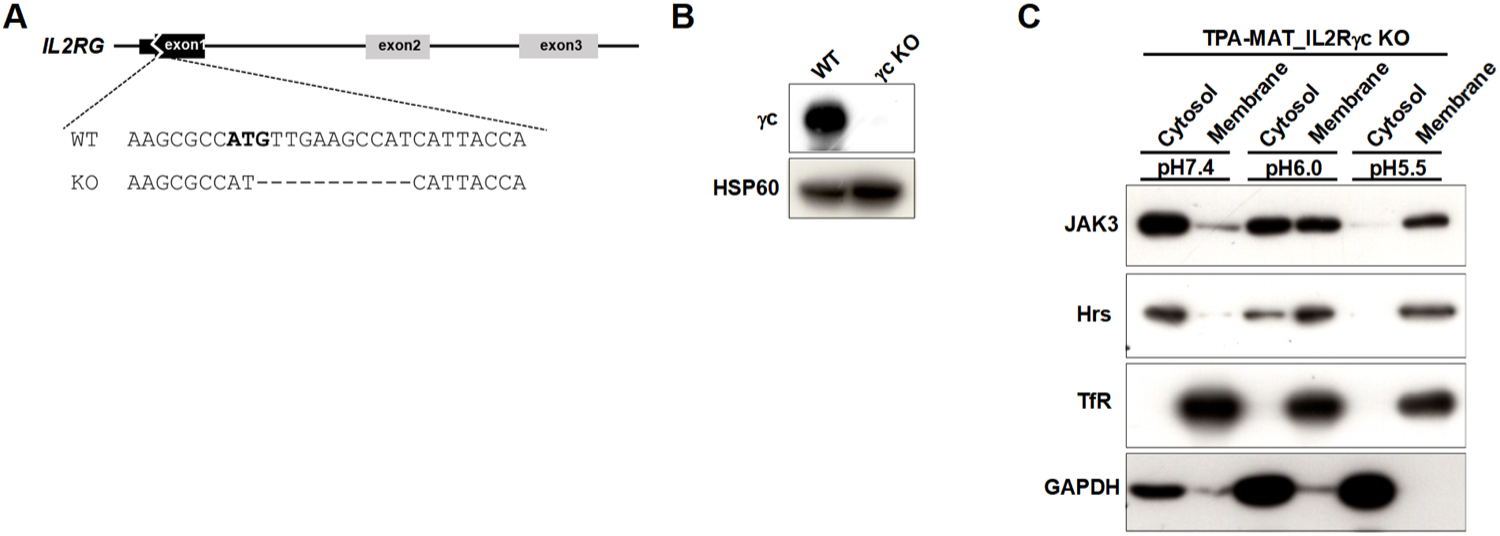
The presence of γc is not required for acidic pH-dependent membrane binding of JAK3. (A) We generated γc knockout TPA-MAT cells using the lentiviral CRISPR/Cas9 system (details are described in the STAR methods). The sequence of the edited region is shown. (B) Confirmation of gene editing was performed by examining protein expression. The expression levels of γc were assessed by western blotting. (C) Acidic pH-dependent membrane interaction of JAK3 in γc knockout TPA-MAT cells. Cells were homogenized at pH 7.4, pH 6.0, and pH 5.5 and fractionated into membrane and cytosol fractions by ultra-centrifugation. GAPDH and Transferrin receptor (TfR) were used as cytoplasmic and membrane fraction markers, respectively.

**Supplementary Figure 3.**
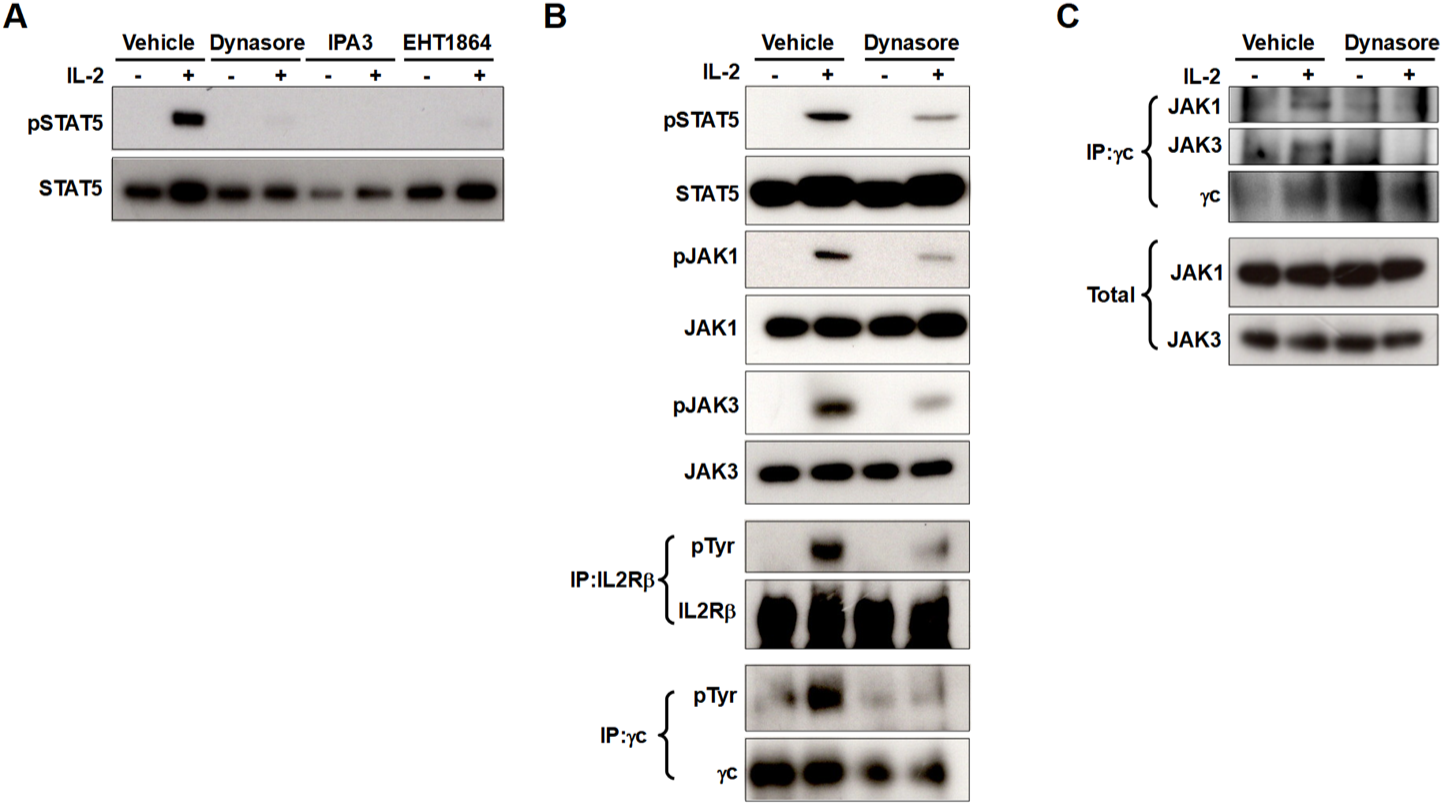
Endocytosis of interleukin (IL)-2 receptor multimer is necessary for functional IL-2 signal complex assembly. (A–C) TPA-MAT cells were pre-incubated with endocytosis inhibitors Dynasore (80 μM), IPA3 (20 μM), or EHT1864 (10 μM), for 30 minutes in serum-free medium and stimulated with 50 ng/mL IL-2 for 10 minutes. The phosphorylation levels of (A) STAT5 and (B) all components of the IL-2 signal complex were assessed by western blotting. (C) The cell lysates were immunoprecipitated with an anti-γc antibody and coprecipitates were analyzed by western blotting. All data are representative of at least three independent experiments.

**Supplementary Figure 4.**
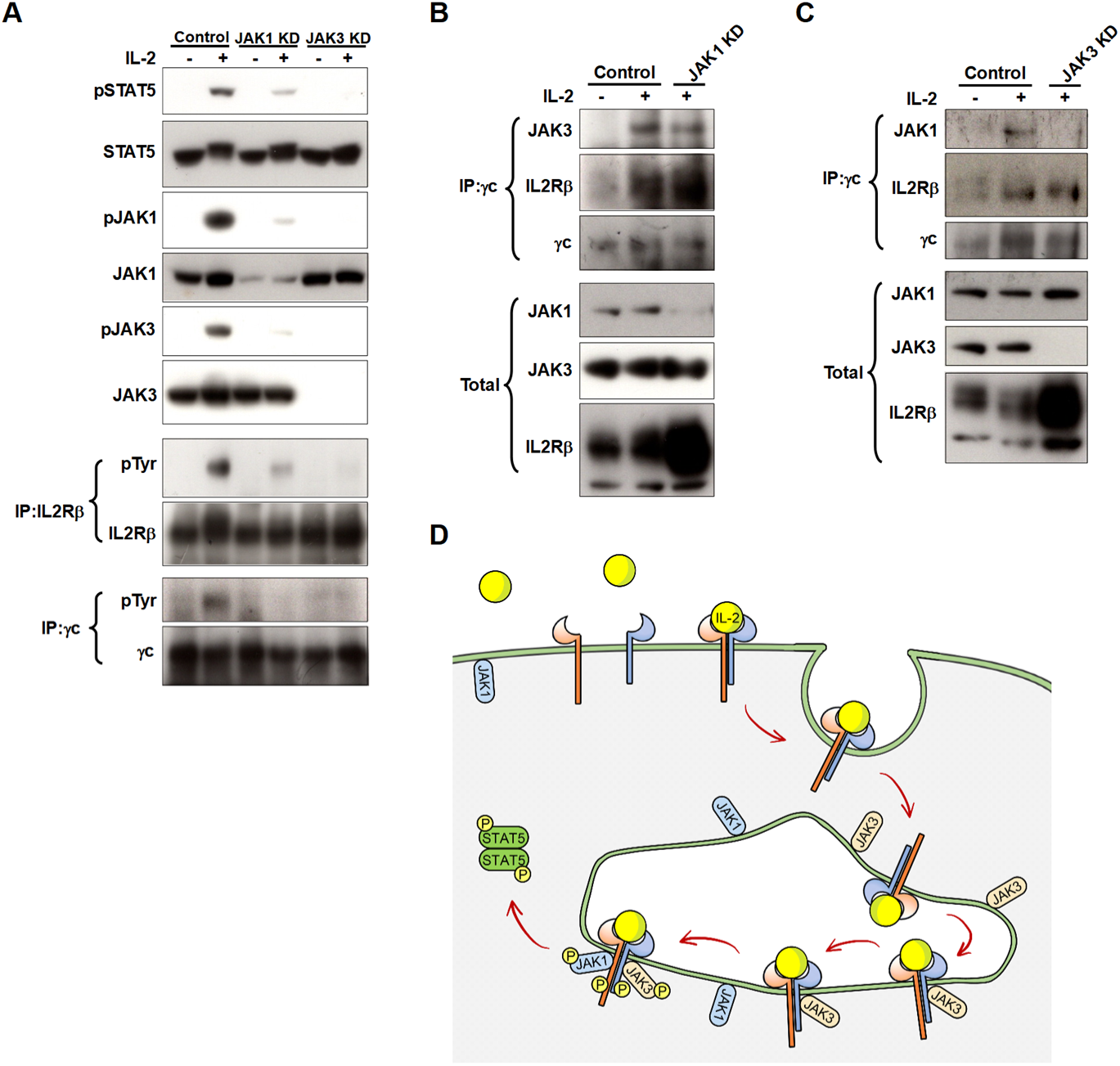
JAK3 is required for the recruitment of JAK1 to the interleukin (IL)-2 signal complex. (A) The effects of JAK1 or JAK3 knockdown on the IL-2 induced signal activation were assessed by western blotting. We performed respective knockdown of JAK1 and JAK3 in TPA-MAT cells using the lentiviral short hairpin RNA (shRNA) expression system (details are described in the STAR methods). Cells were stimulated with IL-2 for 10 minutes. (B, C) Cell lysates from IL-2-stimulated (B) JAK1 knockdown TPA-MAT cells and (C) JAK3 knockdown TPA-MAT cells were immunoprecipitated with an anti-γc antibody and coprecipitates were analyzed by western blotting. (D) Schematic of IL-2-induced signal complex assembly. IL-2 induces receptor multimerization on the plasma membrane, where neither JAK1 nor JAK3 is embedded. Following endocytosis, the IL-2 receptor multimer interacts with JAK3 on endosomes. This step is necessary for the recruitment of JAK1 to the IL-2 signal complex. All components are needed to activate IL-2-induced signal transduction. Data in (A–C) are representative of at least three independent experiments.

